# CELLULOSE SYNTHASE INTERACTIVE1- and Microtubule-Dependent Cell Wall Architecture Is Required for Acid Growth in *Arabidopsis Hypocotyls*

**DOI:** 10.1101/692202

**Authors:** Xiaoran Xin, Lei Lei, Yunzhen Zheng, Tian Zhang, Sai Venkatesh Pingali, Hugh O’Neill, Daniel J. Cosgrove, Shundai Li, Ying Gu

**Affiliations:** Department of Biochemistry and Molecular Biology, Pennsylvania State University, University Park, PA 16802 USA; Department of Biology, Pennsylvania State University, University Park, PA 16802 USA; Biology and Soft Matter Division, Oak Ridge National Laboratory, Oak Ridge, TN 37831 USA

**Keywords:** cell wall, axial elongation, acid growth, microtubules, atomic force microscopy, electron microscopy, crossed-polylamellate walls

## Abstract

Auxin-induced cell elongation relies in part on the acidification of the cell wall, a process known as acid growth that presumably triggers expansin-mediated wall loosening via altered interactions between cellulose microfibrils. Cellulose microfibrils are a major determinant for anisotropic growth and they provide the scaffold for cell wall assembly. Little is known about how acid growth depends on cell wall architecture. To explore the relationship between acid growth-mediated cell elongation and plant cell wall architecture, two mutants (*jia1-1* and *csi1-3*) that are defective in cellulose biosynthesis and cellulose microfibril organization were analyzed. The study revealed that cell elongation is dependent on CSI1-mediated cell wall architecture but not on the overall crystalline cellulose content. We observed a correlation between loss of crossed-polylamellate walls and loss of auxin- and fusicoccin-induced cell growth in *csi1-3*. Furthermore, induced loss of crossed-polylamellate walls via disruption of cortical microtubule mimics the effect of *csi1* in acid growth. We hypothesize that CSI1- and microtubule-dependent crossed-polylamellate walls are required for acid growth in *Arabidopsis* hypocotyls.

## Introduction

Anisotropic cell expansion is a characteristic feature of plant cells. Plant cells respond to developmental and environmental stimuli by altering the growth direction. Cell expansion results from the interplay between uniform turgor pressure and a yielding cell wall. The control of the direction of cell elongation resides in the anisotropic cell wall structure. Cellulose is considered the major load-bearing polymer in the primary cell wall. Therefore, its role in regulating directional cell expansion has been extensively studied. A widely cited hypothesis states that the orientation of cellulose microfibrils is perpendicular to the axis of maximum cell elongation. In other words, transversely oriented cellulose microfibrils in the cell wall favors expansion in the long axis and restricts lateral expansion, an arrangement likened to “hoops around a barrel” (Green, 1960, 1962). However, transverse orientation of cellulose microfibrils is not always correlated with a rapid cell elongation rate. Axially aligned cellulose microfibrils were observed at the innermost cell wall in elongating parenchyma cells (Itoh, 1975; Itoh and Shimaji, 1976). The inhibition of elongation in *procuste*, a cellulose synthase mutant of *Arabidopsis*, was accompanied by normal deposition of transverse orientation of nascent cellulose microfibrils in hypocotyl epidermal cell wall (MacKinnon *et al*., 2006). Similarly, cellulose microfibrils in the innermost cell wall layer were transversely oriented in the elongation zone of *rsw4* and *rsw7* roots in which cell elongation was drastically reduced (Wiedemeier *et al*., 2002). The conflicting data may partly be explained by the differences in regulation in aerial vs. root tissues. Moreover, early studies are limited by failure to observe dynamic reorganization of cellulose microfibrils as inferred by visualization of CESA trajectories (Chan *et al*., 2010). To add another layer of complexity, the alignment of cellulose microfibrils at the inner and outer periclinal walls of epidermal cells differs significantly (Supplementary Fig. S1). Recent discussions have been centered on which cell wall feature is a better predictor of growth anisotropy. Because the outer epidermis has a nearly net isotropic orientation of cellulose, two studies have favored the role of inner periclinal walls of epidermis in the regulation of growth anisotropy (Chan *et al*., 2011; Crowell *et al*., 2011).

However, the outer epidermal cell walls bear most of the stress exerted by the expanding internal tissues and represent a growth-limiting sheath for multicellular systems (Green, 1980; Kutschera, 2008; Savaldi-Goldstein and Chory, 2008). The outer periclinal epidermal walls (~1 μm) of *Arabidopsis* hypocotyl are much thicker than the inner periclinal walls (Derbyshire *et al*., 2007). The inner periclinal walls (~50 nm) are estimated to accommodate at most 1-2 microfibrillar lamellae. In the shoot apical meristem, the outer periclinal epidermal wall is approximately seven times thicker than the inner periclinal wall (Kierzkowski *et al*., 2012). The outer periclinal epidermal walls in onion epidermal cell have ~100 lamellae with different cellulose orientation between each successive lamella (Zhang *et al*., 2016). Consistent with outer periclinal epidermal wall’s role in limiting elongation of the intact organ, studies have shown that outer epidermal-specific expression of the brassinosteroid receptor or a brassinosteroid biosynthetic enzyme in null mutants was sufficient to rescue their dwarf phenotypes in *Arabidopsis*, indicating that molecular components involved in perception of external cues and in transducing into mechanical sensing/wall modification reside in the outer epidermis (Savaldi-Goldstein and Chory, 2008). It remains unclear whether and how the crossed-polylamellate outer epidermal walls regulate cell elongation. It has been postulated that cell elongation is determined by the direction of net alignment among cellulose microfibrils in many cells of a tissue or organ (Baskin, 2005).

Multiple hypotheses have been proposed to explain how the crossed-polylamellate outer epidermal walls are formed. The observations that microfibril orientation changes from transverse to longitudinal from the most recently deposited cell wall to the outermost cell wall of *Nitella* suggest that a passive re-orientation of cellulose microfibrils to axial direction occurs during growth (Gertel and Green, 1977; Green, 1960). As cell wall matrix polysaccharides are continuously deposited during growth, they may contribute to passive re-orientation of microfibrils (Gertel and Green, 1977; Preston, 1982). Alternatively, the anisotropic orientation of microfibril deposition may involve guidance by microtubules (MTs) (Chan *et al*., 2010). The latter concept is supported by the observation that the rotation of the cellulose synthase trajectories was abolished by reagents disrupting microtubules, accompanying the loss of the crossed-polylamellate wall in *Arabidopsis* hypocotyls (Chan *et al*., 2010).

The role of microtubules in guidance of cellulose microfibril alignment is probably not conserved in all cell types, but in general there is good agreement between the orientation of cellulose microfibrils and the underlying cortical microtubules in cells undergo rapid cell elongation (Heath, 1974; Hepler and Palevitz, 1974). Supporting the microtubule-microfibril alignment hypothesis, cellulose synthase complexes (CSCs) have been visualized as diffraction-limited particles that move along the underlying cortical microtubules (Paredez *et al*., 2006). Concurrent rotations have been observed at the outer epidermal cell wall of the hypocotyl for both microtubules and CSC trajectories (Chan *et al*., 2007). CELLULOSE SYNTHASE INTERACTIVE1 (CSI1) is a linker protein that mediates the interaction between CSCs and microtubules (Bringmann *et al*., 2012; Gu *et al*., 2010; Li *et al*., 2012). It does so by interacting directly with both CSCs and microtubules (Li *et al*., 2012). The CSC trajectories were uncoupled from the underlying cortical microtubules in a *csi1* null mutant, supporting its essential role of co-alignment between CSC trajectories and microtubules. As the *CSI1* mutation uncouples CSC trajectories from microtubules, it serves as a tool to re-evaluate the importance of orientation of cellulose microfibrils during rapid cell elongation. An additional useful tool includes *jiaoyao1-1* (*jia1-1*) that bears a missense mutation in KORRIGAN, a membrane-associated endo-β-1,4 glucanase, which results in reduced crystalline cellulose content and decreased cellulose microfibril organization (Lei *et al*., 2014).

In this study, the growth defects in *jia1* and *csi1* mutants were characterized in detail. The results suggest that the alignment of the most recently deposited cellulose microfibrils in the outer periclinal wall is not sufficient to explain anisotropic cell elongation in both mutants. A correlation between loss of crossed-polylamellate wall and loss of auxin- and fusicoccin (FC)-induced cell elongation was revealed.

## Results

### The *csi1-3* mutants display cell expansion defects

To understand the relationship between cellulose microfibril organization and cell elongation, we used *jia1-1* and *csi1-3* mutants that disrupt cellulose biosynthesis. Dark-grown *Arabidopsis* hypocotyls are a widely used model system to study cell growth (Derbyshire *et al*., 2007; Gendreau *et al*., 1997). *csi1-3* was previously shown to have reduced elongation by 40% in dark-grown hypocotyls and the reduction of growth rate was maximal at 3 to 5 days after cold stratification (Gu *et al*., 2010). We compared growth morphology of four-day-old dark-grown hypocotyls in the two mutants. Both *csi1-3* and *jia1-1* exhibited dwarf hypocotyls (Supplementary Fig. S2A). Similar to wild type hypocotyls, both *csi1-3* and *jia1-1* had approximately 22 (± 2) epidermal cells from base to the apical hook (Supplementary Fig. S2C). As growth in dark-grown hypocotyls does not involve cell division, the reduced hypocotyl length reflects a reduced length of epidermal cells. In dark grown hypocotyls, epidermal cells elongate in an acropetal fashion in which growth is initiated in the base of the hypocotyl and the elongation zone moves up with time (Gendreau *et al*., 1997, Supplementary Fig. 2C). Compared with wild type (WT), *jia1-1* mutants had reduced cell length in cells #10 to #24 (Supplementary Fig. S2D). By day 10 after germination, dark grown hypocotyls reach their maximum lengths (Gendreau *et al*., 1997; Gu *et al*., 2010). WT and *jia1-1* hypocotyls had approximately similar lengths by 10-day post germination, indicating that *jia1-1* hypocotyls had a delay in acropetal growth acceleration and eventually caught up with wild-type hypocotyls. 4-day-old *csi1-3* hypocotyls elongated less than WT hypocotyls and they did not catch up by 10-day post germination (Supplementary Fig. S2B). These data suggest that reduced cell elongation in *csi1-3* appeared to be independent of its position or time.

### The *csi1-3* mutants are deficient in auxin- or FC-induced hypocotyl elongation

In the classic acid growth hypothesis, cell elongation is induced by acidification of thick extension-limiting outer epidermal cell walls, which can be achieved by either the growth hormone auxin (indole-3-acetic acid, IAA) or the phytotoxin fusicoccin (Hager *et al*., 1971; Marre, 1979; Rayle and Cleland, 1970; Rayle, 1973; Rayle and Cleland, 1977; Rayle and Cleland, 1992). Both IAA and FC induce decreases in apoplastic pH, which in turn activate wall-loosening agents such as expansin, resulting in cell wall modification/expansion (Cosgrove, 1997, 2000; Kutschera, 1994; McQueen-Mason *et al*., 1992). As an approach to understand the control of axial cell elongation in living cells, we adopted a hypocotyl segment system to test how *csi1-3* and *jia1-1* respond to exogenous IAA (Cosgrove *et al*., 2002; Fendrych *et al*., 2016). To deplete endogenous auxin, shoot apexes and roots were removed from 3-day-old etiolated *Arabidopsis* seedlings. To find a minimum IAA concentration that induces sufficient hypocotyl segment elongation, we tested a series of exogenous IAA concentrations and chose 1 μM of IAA for the following experiments (Supplementary Fig. S3A). It has been reported that auxin induced hypocotyl segment elongation occurred after approximately 20 min of auxin application and lasted approximately 2 hours (Fendrych *et al*., 2016). Thus, we recorded the final length of hypocotyl segment at 150 min after the application of IAA. The hypocotyl segment corresponds to cells #5-17. The auxin-induced hypocotyl segment elongation was ~ 60% of the elongation rate of intact hypocotyls under normal growth conditions (Supplementary Fig. S4). The results showed that exogenous auxin induced similar elongation of the hypocotyl segment of wild type and the *jia1-1* mutants while hypocotyl elongation was impaired in the *csi1-3* mutants (Fig. 1A and Supplementary Fig. S4B). The elongation rate for auxin-induced hypocotyl segment was similar for wild type (96.6 ± 7.5 µm/h) and *jia1-1* (96.3 ± 17.1 µm/h) but significantly reduced in *csi1-3* (22.5 ± 9.8 µm/h).

**Fig. 1.**
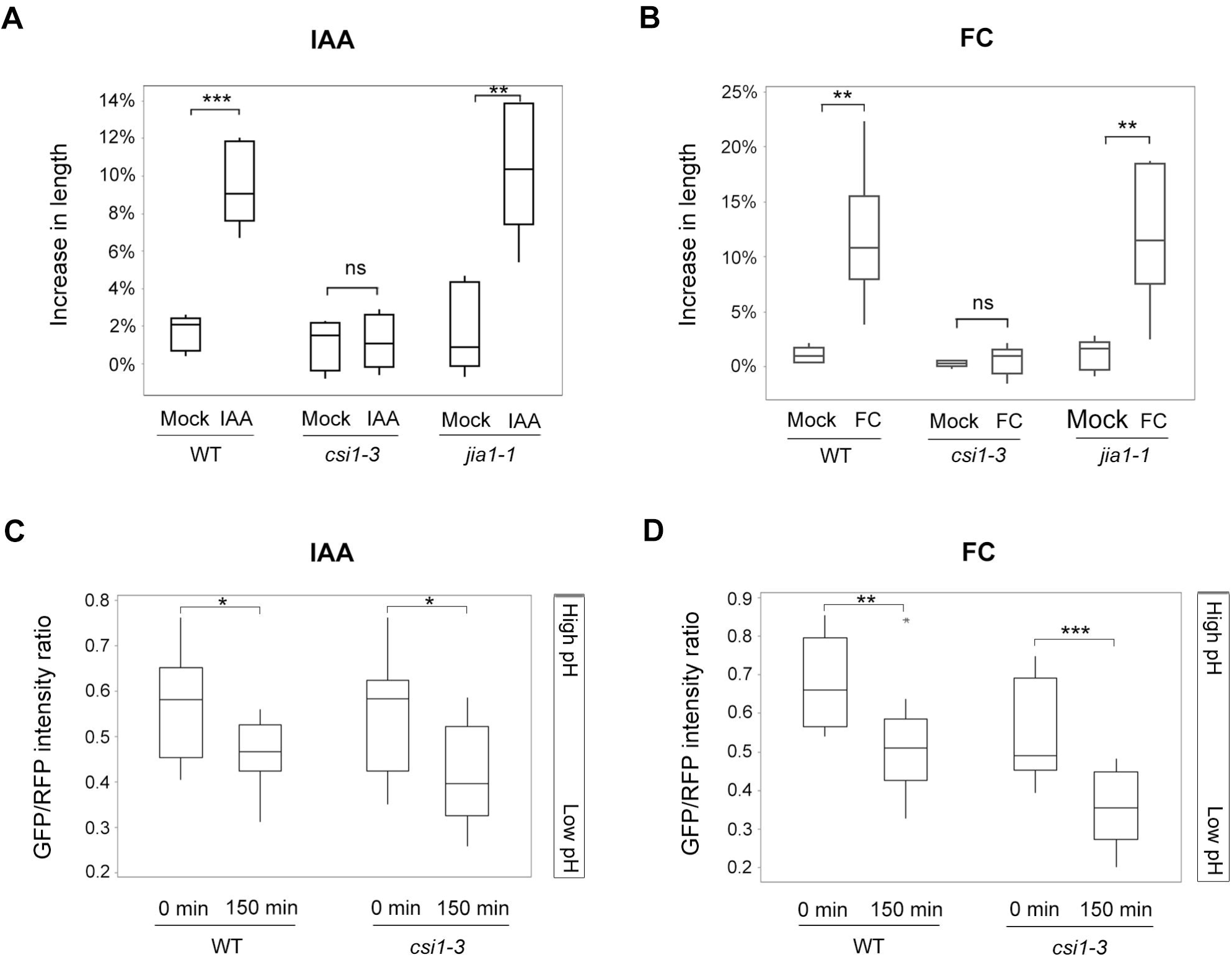
The *csi1-3* mutants are not responsive to auxin- or FC-induced hypocotyl elongation. (A) Auxin-induced hypocotyl segments elongation of wild type (Col-0), *csi1-3*, and *jia1-1* mutants. One out of three reproducible independent experiments were used in the analysis. The lengths of the hypocotyls were measured at 0 min and 150 min of each treatment. ** *P* ≤ 0.01, *** *P* ≤ 0.001 (n = 5 in mock treatment for each genotype, n = 6 in IAA treatment for each genotype). Statistical analysis was performed by two-tailed Student’s *t*-test. (B) FC-induced hypocotyl segments elongation of wild type (Col-0), *csi1-3*, and *jia1-1* mutants. One out of three reproducible independent experiments were used in the analysis. The lengths of the hypocotyls were measured at 0 min and 150 min of each treatment. ** *P* ≤ 0.01 (n = 5 in mock treatment for each genotype, n = 6 in FC treatment for each genotype). Statistical analysis was performed by two-tailed Student’s *t*-test. (C) Apoplastic pH change of wild type (Col-0) and *csi1-3* in response to IAA treatment. Images were taken at 0 min and 150 min of IAA treatment. The apoplastic pH was reflected by GFP/RFP intensity ratio. * *P* ≤ 0.05 (n = 12 for each genotype). Statistical analysis was performed by two-tailed Student’s *t*-test. (D) Apoplastic pH change of wild type (Col-0) and *csi1-3* in response to FC treatment. Images were taken at 0 min and 150 min of FC treatment. The apoplastic pH was reflected by GFP/RFP intensity ratio. ** *P* ≤ 0.01, *** *P* ≤ 0.001 (n = 12 for each genotype). Statistical analysis was performed by two-tailed Student’s *t*-test.

To further verify that *csi1* was not responsive to acid-induced growth, we used the fungal toxin fusicoccin, which activates the plasma membrane (PM) H^+^-ATPase independent of the auxin-mediated mechanism (Baunsgaard *et al*., 1998; Rayle and Cleland, 1992). We found that 2 μM of FC was sufficient to trigger the growth of wild type hypocotyl at comparable levels to that of 1 μM IAA (Supplementary Fig. S3B). Thus, this concentration was used in the following assays. The hypocotyl segment growth triggered by FC has been observed to occur continuously within 2 hours (Fendrych *et al*., 2016). Therefore, we chose to measure the final length of hypocotyl 150 min after FC treatment. *csi1-3* mutants were not responsive to fusicoccin-induced hypocotyl elongation, whereas wild type and *jia1-1* mutants exhibited normal responses (Fig. 1B).

### Cell wall acidification is normal in the *csi1-3* mutants

To examine whether cell wall acidification occurs normally in the *csi1-3* mutants, we used a classic method to measure the apoplastic pH of *Arabidopsis* etiolated seedlings. The pH of the incubation medium was measured, from which cell wall acidification can be inferred (Cleland, 1973). We used approximately thirty 4-day-old etiolated seedlings in each measurement. After 150 min of 1 μM IAA incubation, the pH of the incubation medium dropped from 6.05 (± 0.02) to 5.88 (± 0.08) in wild type and from 6.05 (± 0.02) to 5.87 (± 0.07) in *csi1-3* mutants, indicating that cell wall acidification is not impaired in the *csi1-3* mutants.

The apoplastic pH sensor apo-pHusion was recently developed to allow direct measuring apoplastic pH changes in epidermal cells (Fendrych *et al*., 2016; Gjetting *et al*., 2012). We crossed *csi1-3* with the line containing the apo-pHusion sensor, which was a tandem fusion of two fluorescent proteins, mRFP1 and EGFP. A chitinase targeting sequence was added in the construct to target the sensor to the apoplast (Gjetting *et al*., 2012). The mRFP1 is considered to be insensitive to pH changes in the physiologically relevant range, whereas EGFP is sensitive to pH changes and low pH inhibits the EGFP fluorescence (Gjetting *et al*., 2012). Images of both RFP and GFP channels were taken at 0 min and 150 min of IAA and FC treatment. The ratio of GFP/RFP fluorescent intensity was measured and calculated in Image J, where RFP intensity was used as an intramolecular reference. The results showed that in both wild type and *csi1-3* mutants the GFP/RFP intensity ratio dropped after IAA or FC treatment, indicating a drop in apoplastic pH and normal apoplastic acidification (Fig. 1C and 1D). The normal acidification in *csi1-3* indicates that the loss of auxin-induced cell elongation in *csi1-3* is not due to deficient apoplastic acidification.

### The *csi1-3* mutants have less extensible cell walls

The cell wall creep assay has been used to mimic the slow creep of cell walls during cell expansion (Cosgrove, 1998; Durachko and Cosgrove, 2009; Park and Cosgrove, 2012b). We measured the acid-induced creep responses of native walls. To kill the cells, 3-day-old etiolated seedlings were frozen and thawed prior to the experiment. The specimens were clamped at constant force (2.5 g) in pH 6.8 buffer. After 20 min incubation, the buffer was replaced with pH 4.5 buffer. The walls of wild type and *jia1-1* mutants extended rapidly in response to acidic buffer. In contrast, the wall extension and extension rate of the *csi1-3* mutants was approximately 41% and 24% of wild-type hypocotyls, respectively (Fig. 2A and 2B). Although both *csi1-3* and *jia1-1* had reduced cell elongation in 3-day-old de-etiolated hypocotyls *in vivo*, only *csi1-3* had diminished creep response to acidic buffers. These results support the idea that different mechanisms account for cell expansion defects in the two mutants.

**Fig. 2.**
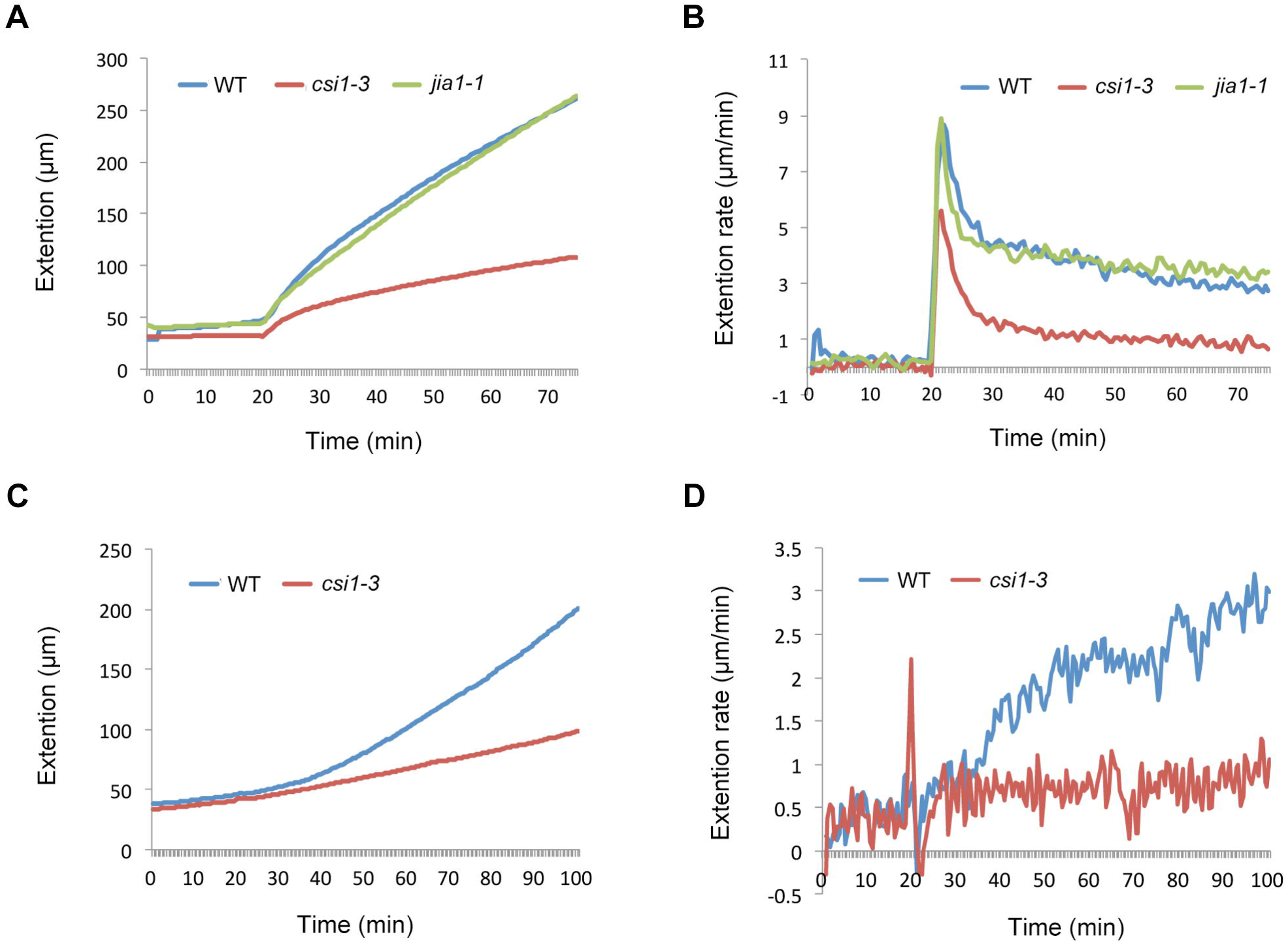
The *csi1-3* mutants exhibit less wall extensibility. (A and B) Extension of native walls in response to acidic buffer (pH 4.5). The samples were 3-day-old dark-grown seedlings of wild type (Col-0), *csi1-3*, and *jia1-1* mutants. Each curve is an average of eight individual responses. (C and D) Extension of heat-inactivated wild type and *csi1-3* walls in pH 4.5 buffer upon the addition of 50 μg/mL GH5 family endoglucanase at 20 min. The samples were leaf petioles from wild type (Col-0) and *csi1-3* mutant. Each curve is an average of at least four individual responses (n = 4 for wild type, n = 5 for *csi1-3*).

We next tested whether *csi1* affects an endoglucanase-induced creep response as a means to test the wall’s ability to undergo creep. It was shown previously that a creep response was induced by PpXG5, a family-5 endoglucanase (Park and Cosgrove, 2012b). The GH5 family endoglucanase displays hydrolytic activity towards both xyloglucan and disordered cellulose (Park and Cosgrove, 2012b). To inactivate endogenous enzymes and expansins, samples were boiled for 12 s and immediately cooled down with cold running water. Because *Arabidopsis* hypocotyls were not amenable to heat inactivation followed by creep experiments, we used leaf petiole instead. The leaf petioles were boiled to inactivate endogenous wall-loosening enzymes and abraded to promote enzyme penetration through the cuticle. The wall specimens were clamped at constant force (8.2 g) and 50 μg/mL GH5 family endoglucanase was added after 20 min of incubation. The *csi1-3* samples showed a small creep rate compared to that in wild type (Fig. 2D). Diminished creep responses in *csi1* suggest that the reduced wall extensibility in *csi1-3* is likely due to changes in wall architecture.

### The *csi1-3* mutants display altered cellulose microfibril organization

It is commonly accepted that the direction of maximal expansion depends on the direction of the alignment of cellulose microfibrils. To assess whether the reduced cell elongation in *csi1-3* and *jia1-1* is correlated with mis-alignment of cellulose microfibrils, we examined the organization of cellulose microfibrils. Atomic force microscopy (AFM) was used to examine the organization of the most recently deposited cellulose microfibrils in outer periclinal walls in epidermal cells of *Arabidopsis* hypocotyls (Supplementary Fig. S1). The hypocotyl samples were kept in water during sample preparation and imaging, so the organization of cellulose microfibrils was maintained in a near-native state (Zhang *et al*., 2016). AFM images revealed that the majority of most recently deposited cellulose microfibrils in the outer periclinal walls were arranged in transverse orientation perpendicular to the growth axis in wild type seedlings (Fig. 3A and 3C). Although the majority of cellulose microfibrils were deposited in transverse orientation in *csi1-3* mutants, there was less variation in orientation of cellulose microfibrils as compared with wild type (Fig. 3A and 3C). Similar parallel alignment of cellulose microfibrils was previously shown for *jia1-1* (Lei *et al*., 2014). *csi1-3* and *jia1-1* mutants exhibited different cell expansion abilities but similar parrallel alignment of the most recently deposited cellulose microfibrils, indicating that cellulose microfibril alignment in the innermost outer periclinal wall alone is insufficient to regulate axial cell elongation.

**Fig. 3.**
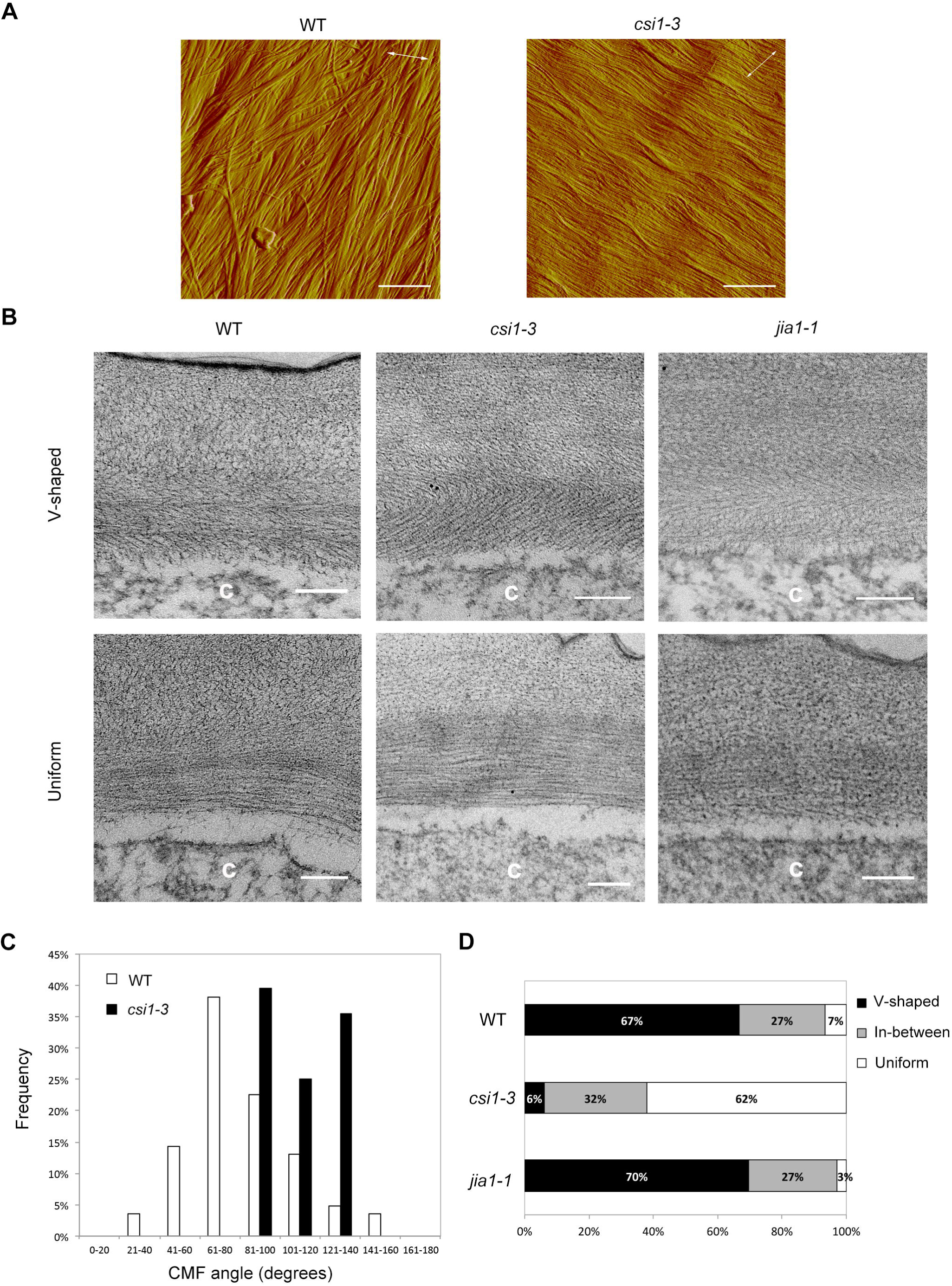
Cellulose microfibril organization is altered in the *csi1-3* mutants. (A) AFM images of 3-day-old dark-grown seedlings of wild type (Col-0) and *csi1-3* mutant. The long axes of the cells are indicated by white double-headed arrows. (Scale bars, 400 nm.) (B) TEM images of 3-day-old dark-grown seedlings of wild type (Col-0), *csi1-3*, and *jia1-1* mutants. (Scale bars, 200 nm.) The letter “c” on the TEM images indicates cytosol of the cell. (C) Quantitative analysis of cellulose microfibril (CMF) angles in AFM images. n = 84 for wild type, n = 48 for *csi1-3*. (D) Quantitative analysis of the TEM images. n = 30 for each genotype. V-shaped refers to ‘herringbone’ pattern of cell wall. Uniform refers to unifying parallel orientation of polysaccharides. In-between is a category that has mixed features of V-shaped and uniform cell wall.

### The crossed-polylamellate architecture of outer epidermal cell wall is lost in the *csi1-3* mutants

It has been hypothesized that the outer epidermal wall represents the growth-limiting structure of the multicellular system in hypocotyls (Kutschera, 1992). We therefore examined the structure of the outer epidermal wall using transmission electron microscopy (TEM). Polysaccharides in oblique sections of 3-day-old *Arabidopsis* etiolated hypocotyls 0.5-2 mm below the apical hook were stained with periodic acid thiocarbohydrazide-silver proteinate (PATA) and lead citrate. In wild type cells, epidermal cell walls have a crossed-polylamellate architecture in which cellulose microfibrils are relatively well aligned within each lamella but the orientation varies between successive lamellae (Fig. 3B and 3D). Occasionally outer epidermal walls show uniform parallel orientation of polysaccharides across multiple lamellae in wild type (7%). A comparable texture was observed for *jia1-1* (Fig. 3B and 3D). In contrast, majority of outer epidermal walls (62%) showed uniform parallel orientation of polysaccharides in different lamellae in *csi1-3* (Fig. 3B and 3D).

### Pharmacological disruption of MTs phenocopies loss of auxin- and FC-induced cell expansion and loss of the crossed-polylamellate wall architecture in *csi1-3*

CSI1 was previously shown to be required for the co-alignment between CSC trajectories and cortical microtubules (Gu *et al*., 2010; Li *et al*., 2012). In *csi1-3*, CSC trajectories did not follow the underlying microtubules in epidermal cells approximately 0.5 mm below the apical hook (Li *et al*., 2012). As the cell expansion defect is not limited to the upper part of hypocotyl, we carefully examined dynamics of YFP-CESA6 and cortical microtubules in all regions of hypocotyls. In wild type seedlings, microtubule orientation and CSC trajectories were consistently aligned across different regions, from predominantly transversely oriented in cell #17-21, to oblique orientation in cell #9-13, to longitudinal orientation in cell #1-5. In contrast, *csi1-3* seedlings displayed consistently transverse orientation of CSC trajectories in all regions, while microtubule orientation were similar to that of wild type seedlings (Supplementary Fig. S5). The angles of MT or CSC tracks against horizontal direction were measured using FibrilTool, an Image J plug-in to quantify fibrillar structures in microscopy images (Boudaoud *et al*., 2014), further supporting the conclusion that the average trajectories of YFP-CESA6 and MTs were uncoupled in *csi1-3* (Li *et al*., 2012, Supplementary Fig. S5).

The loss of crossed-polylamellate walls in *csi1-3* is consistent with the hypothesis that crossed-polylamellate walls of the outer epidermal cell require microtubule-dependent rotation of cellulose microfibril deposition (Chan *et al*., 2007; Chan *et al*., 2010). To further explore microtubules’ role in cell expansion, we treated wild type seedlings with oryzalin, a MT depolymerization drug. We chose an oryzalin concentration that mimicked the morphology of *csi1-3* in terms of the cell expansion defects (Li *et al*., 2012). Oryzalin-treated hypocotyls did not elongate in response to auxin or FC treatments, thus mimicking the response of *csi1-3* (Fig. 4A and 4B). Moreover, TEM images of oryzalin-treated seedlings also showed a loss of the crossed-polylamellate wall architecture (Fig. 4C and 4D). The majority of oryzalin-treated seedlings displayed a uniform parallel orientation of polysaccharides (24/30 images). These results suggest that MTs play an important role in the formation of the crossed-polylamellate wall architecture and loss of auxin-induced cell elongation accompanies loss of crossed-polylamellate construction.

**Fig. 4.**
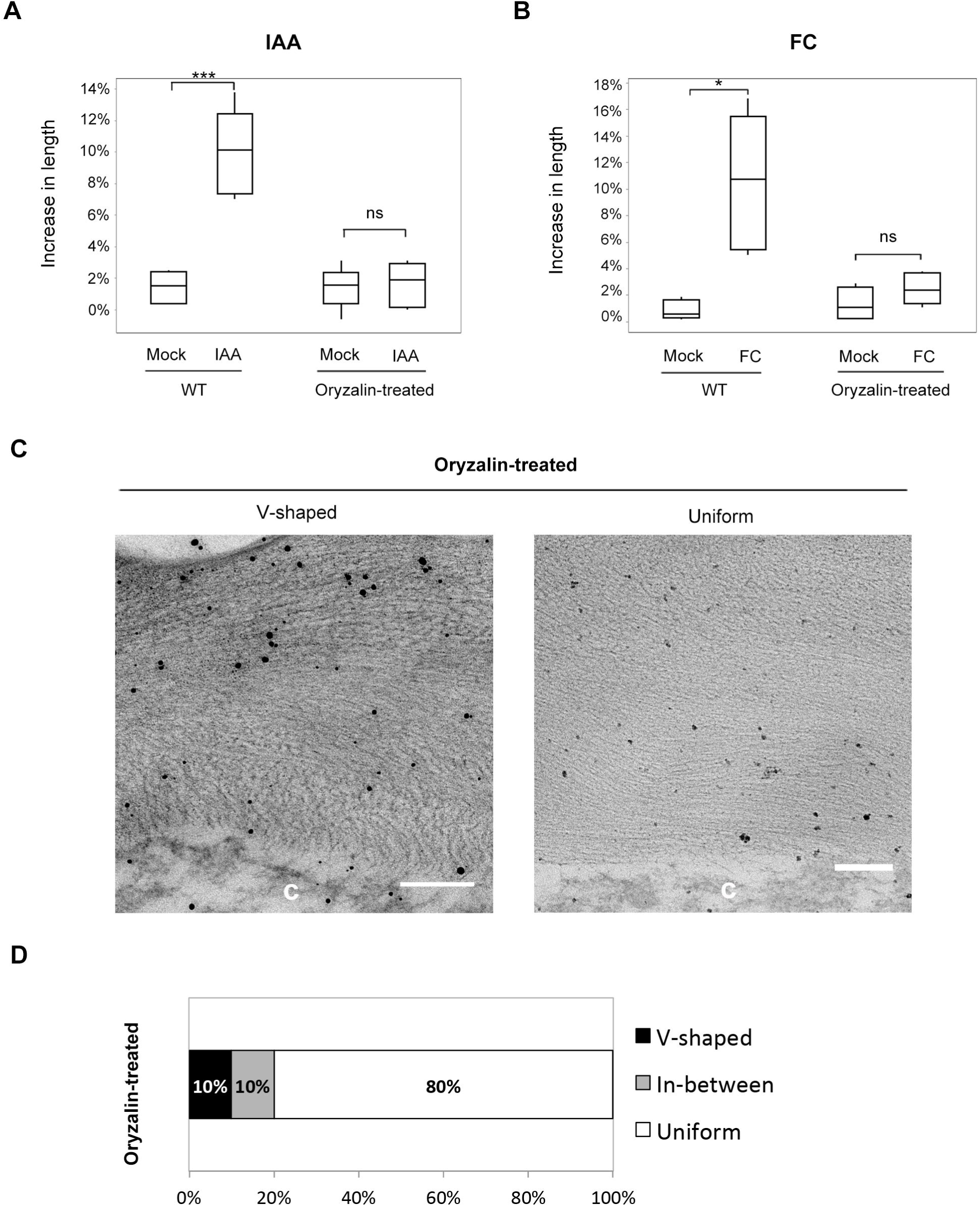
Pharmaceutical disruption of microtubules results in loss of the crossed-polylamellate wall architecture. (A) Auxin-induced hypocotyl segments elongation of wild type (Col-0) treated with 300 nM oryzalin. One out of three reproducible independent experiments were used in the analysis. The lengths of the hypocotyls were measured at 0 min and 150 min of each treatment. *** *P* ≤ 0.001 (n = 6 in each treatment for both wild type and oryzalin-treated seedlings). Statistical analysis was performed by two-tailed Student’s *t*-test. (B) FC-induced hypocotyl segments elongation of wild type (Col-0) treated with 300 nM oryzalin. One out of three reproducible independent experiments were used in the analysis. The lengths of the hypocotyls were measured at 0 min and 150 min of each treatment. * *P* ≤ 0.05 (n = 5 in each treatment for both wild type and oryzalin-treated seedlings). Statistical analysis was performed by two-tailed Student’s *t*-test. (C) TEM images of 3-day-old dark-grown seedlings of wild type (Col-0) treated with 300 nM oryzalin. (Scale bars, 200 nm.) The letter “c” on the TEM images indicates cytosol of the cell. (D) Quantitative analysis of the TEM images. n = 30. V-shaped refers to ‘herringbone’ pattern of cell wall. Uniform refers to unifying parallel orientation of polysaccharides. In-between is a category that has mixed features of V-shaped and uniform cell wall.

## Discussion

### Crossed-polylamellate cell wall architecture

Crossed-polylamellate walls are often observed in rapidly growing epidermal cells in both hypocotyls and roots (Chafe and Wardrop, 1972; Hodick and Kutschera, 1992; Roland *et al*., 1977; Takeda and Shibaoka, 1981). It is postulated that different mechanisms account for crossed-polylamellate wall feature in two systems (Li *et al*., 2014). In *Arabidopsis* hypocotyls, cyclic reorientation of microtubules is accompanied with a corresponding reorientation of CSC trajectories. The dynamics of cortical microtubule pattern actively dictates orientation of cellulose microfibril deposition. The rotation of CSC trajectories was blocked by either stabilizing or depolymerizing microtubules, and this resulted in loss of crossed-polylamellate cell wall patterns (Chan *et al*., 2010). The cyclic reorientation appears to be shoot-specific as microtubules do not rotate in roots in *Arabidopsis*. In *Arabidopsis* roots, there is a one-way microfibril reorientation from transverse to longitudinal as epidermal cells elongate (Anderson *et al*., 2010; Granger and Cyr, 2001; Liang *et al*., 1996; Sugimoto *et al*., 2000). The one-way microfibril reorientation is most likely a passive process independent of microtubules. Consistent with the idea that shoot and root adopt different mechanisms in growth controls, auxin promotes cell expansion in aerial tissues but inhibits cell expansion in roots at concentrations above 10^−8^ M (Ishikawa and Evans, 1993; Taiz, 1984). *csi1* resulted in the loss of CSC trajectory re-orientation and loss of crossed-polylamellate cell wall patterns, supporting the idea of a microtubule-dependent mechanism in hypocotyls (Lloyd, 2011). Further supporting the microtubule-dependent mechanism, the removal of microtubules by oryzalin phenocopies the loss of crossed-polylamellate cell wall patterns in *csi1*.

### Correlation between loss of crossed-polylamellate wall architecture and acid-induced growth

Auxin-induced growth is characterized by activation of plasma membrane proton pumps that leads to acidification of the cell wall. The acidification of apoplastic space activates expansin that loosens the connection between cellulose microfibrils and non-cellulosic polysaccharides. As a result, cell expansion occurs. *csi1* has defects in auxin- or FC-induced growth while the apoplastic acidification is normal, suggesting the defect may reside in wall loosening. Indeed, *csi1* walls are less extensible in creep assays. As wall creep mimics wall enlargement during cell growth, these results confirm a correlation between defects in wall mechanics and loss of acid-induced growth in *csi1*. Because pharmacological disruption of MTs phenocopies loss of auxin- and FC-induced cell expansion and loss of the crossed-polylamellate wall architecture in *csi1*, we reasoned that MTs play an important role in the formation of the crossed-polylamellate wall and are dependent on CSI1. In addition, the correlation between loss of crossed-polylamellate wall and loss of auxin- and FC-induced cell growth relies on both MT and CSI1 function.

Previous results show that xyloglucan-deficient mutants *xxt1 xxt2* had diminished responses to FC-induced growth and diminished creep responses to acid buffers (Park and Cosgrove, 2012a). Interestingly, the wall architecture is similar to that of *csi1*, both displaying well aligned transversely oriented cellulose microfibrils in the innermost wall and partial loss of crossed-polylamellate wall (Xiao *et al*., 2016). Xyloglucan is speculated to reduce interactions between cellulose microfibrils and reduce their bundling (Xiao *et al*., 2016). It is unclear whether the defects in wall mechanics and/or loss of acid-induced growth in *xxt1 xxt2* alters MT organization or CSI1 function. Transmission electron microscopy analysis revealed an overall appearance of loss of crossed-polylamellate wall in *xxt1 xxt2* and *csi1* but it cannot discern molecular difference such as bundle sizes, microfibril length, connection between different lamellae, microfibril motions, and the incorporation of other cell wall polymers. Precise measurements are required to determine how such cell wall parameters affect the overall wall organization and wall mechanical properties in *xxt1 xxt2* and *csi1*. The mechanism by which cell wall architecture affects acid growth is unclear. Potential mechanisms may involve the inaccessibility of expansin to specific wall-loosening sites, changes in interconnectivity between cellulose microfibrils and other cell wall polymers, and defects in the mechanism of cellulose microfibril sliding or slippage (Cosgrove, 2018).

### Is organization of cellulose microfibrils dependent on microtubules?

The “direct guidance model” postulates that microtubules guide the deposition of cellulose microfibrils (Heath, 1974). The observation that the CSCs trajectories were uncoupled from microtubules in *csi1* supports the direct guidance model. With the discovery of the linker protein CSI1, a modified guidance model is proposed in which microtubules guide the deposition of cellulose microfibrils through direct attachment mediated by CSI1 (Gu *et al*., 2010; Li *et al*., 2012). As CSC trajectories are proxies for the orientation of nascent cellulose microfibrils, it is predicted that cellulose microfibrils would have disorganized orientations in *csi1*. Surprisingly, the newly deposited cellulose microfibrils in *csi1* mutants are more parallelly aligned than those of wild type. These results suggest that the direct guidance model is inadequate to explain the origin of the initial orientation of cellulose microfibrils. A re-visit of alternative models is needed. The self-assembly mechanism suggests that wall formation is a spontaneous process, similar to liquid crystal formation (Neville, 1985; Neville *et al*., 1976). Another model takes the density of CSC and geometry of the cell into consideration (Dyson and Jensen, 2010; Emons and Mulder, 2000). The helicoidal wall texture in root hair was simulated based on the geometrical theory. As *Arabidopsis* hypocotyls have a crossed-polylamellate wall rather than a helicoidal wall, it remains to be tested whether the geometrical theory holds true in hypocotyls.

### Is microfibril alignment sufficient to explain axial cell elongation?

The observations that microtubules and microfibrils in the most recently deposited walls had a dominant orientation perpendicular to the growth axis led to the hypothesis that inner periclinal walls control the growth direction whereas the outer periclinal walls determine the rate of organ elongation (Crowell *et al*., 2010; Crowell *et al*., 2011; Kutschera, 2008). Despite the differences in response to auxin- and FC-induced acid growth in *csi1* and *jia1*, elongation still occurs in both mutants, consistent with the proposed role of inner periclinal walls controlling the growth direction. However, the most recently deposited cellulose microfibrils in *csi1* and *jia1* were more parallelly aligned than those in wild type plants. Therefore, the growth defect in *csi1* and *jia1* cannot be simply explained by the disorganization of the microfibrils and the loss of growth anisotropy. Moreover, the growth defect in *csi1* and *jia1* appeared to be controlled by different mechanisms as *csi1* had reduced cell elongation independent of the cell position along hypocotyls whereas *jia1* had developmental delays in acropetal growth acceleration but eventually was able to reach maximal growth similar to wild type. The synthesis of cellulose is thought to occur simultaneously with elongation to allow continuous deposition of cellulose microfibrils in the innermost layer. Previous experiments have shown that cellulose deposition and net wall polysaccharide synthesis are not tightly linked to the rate of the elongation (Derbyshire *et al*., 2007; Refregier *et al*., 2004). *jia1* had a much more severe defect in cellulose synthesis as compared with *csi1* (Lei *et al*., 2014). It is likely that the delay in growth acceleration represents a mechanism to cope with severe cellulose deficiency.

In summary, this study has several significant implications. First, CSI1 is required for the formation of crossed-polylamellate wall as loss of CSI1 resulted in loss of crossed-polylamellate walls. Second, the crossed-polylamellate wall architecture is required for auxin-induced acid growth. Third, cellulose microfibrils maintain a default transverse orientation without CSI1, indicating the orientation of nascent cellulose microfibrils is independent of CESA-MT linkage. Last, microfibril alignment is not sufficient to control axial cell elongation. Our results lay a foundation for future studies on mechanistic role of wall architecture during acid growth.

## Materials and Methods

### Plant Materials and Growth Conditions

All seeds were surface sterilized with 30% (vol/vol) bleach for 15 min, thoroughly washed with autoclaved double-distilled H_2_O (ddH_2_O), and stored at 4 °C for a minimum of 3 d. Dark-grown seedlings were grown on vertical half-strength Murashige and Skoog (MS) plates without sucrose at 21 °C in the dark. Light-grown seedlings were grown on vertical half-strength MS plates with 1% sucrose at 21 °C on a 16-h light/8-h dark cycle.

### Live-Cell Imaging and Analysis

Images were obtained from epidermal cells of 3-d-old etiolated seedlings 0.5 −2 mm below the apical hook unless indicated otherwise. Imaging was performed on a Yokogawa CSUX1 spinning-disk system as previously described (Li *et al*., 2012). Image analysis was performed using Metamorph and ImageJ software. The average trajectory orientation was analyzed using FibrilTool, an ImageJ plug-in to quantify fibrillar structures (Boudaoud *et al*., 2014).

### Atomic force microscopy

The *Arabidopsis* hypocotyl sample preparation and atomic force microscopy of hypocotyl walls were performed as previously described (Lei *et al*., 2014). The cell angle information was recorded during image acquisition. The microfibril angles were measured using FibrilTool, an ImageJ plug-in to quantify fibrillar structures (Boudaoud *et al*., 2014). Three regions of interest (ROI) were randomly selected from each AFM image for angle measurement.

### Transmission electron microscopy

The apical portions of hypocotyls were cut approximately 0.5 - 2 mm below the apical hook of 3-d-old dark-grown wild type (Col-0), *csi1-3*, and *jia1-1* seedlings. Samples were fixed in 50 mM phosphate buffer (pH 7.0) containing 2.5% glutaraldehyde and 0.1% Triton X-100 for 3 h at room temperature. The residual glutaraldehyde was removed by washing the samples with 50 mM phosphate buffer 3 times for 5 min each. Samples were then immersed in dimethyl sulfoxide (DMSO) for 16 h to cause cell wall swelling and washed with double distilled water (ddH_2_O), followed by dehydration with an ethanol series (25%, 50%, 60%, 70%, 80%, 90%, 100%; 10 min for each); 100% EM grade ethanol 3 times for 10 min; 100% acetone 3 times for 10 min. The dehydrated samples were infiltrated with acetone: Eponate 12 resin (2:1, 1:1, 1:2, > 6 h each) and pure Eponate 12 (3 times for > 6 h). Samples were then embedded in a mold, in which the hypocotyl hook was positioned on the top of mold. After incubating at 70°C overnight, ultrathin sections (~70 nm) were cut at room temperature using an ultra-microtome (Leica EM UC6) equipped with a diamond knife and collected with gold grids.

The thin sections on the grids were incubated with 1% periodic acid in deionized water for 30 min and washed with deionized water 3 times before being incubated with freshly prepared 0.2% thiocarbohydrazide in 20% acetic acid for 24 h. Then the sections were washed sequentially with 10%, 5%, 2% acetic acid, and deionized water. The samples were stained with 1% silver proteinate in deionized water for 30 min and followed by washing. The sections were stained with 0.46% lead citrate in deionized water for 12 minutes, then washed by 25 mM NaOH, 12.5 mM NaOH and deionized water for 1 min each. Images were collected using a transmission electron microscopy (FEI Tecnai TEM) under the 120-kV accelerating voltage conditions.

### Environmental scanning electron microscopy

Etiolated seedlings were gently attached to sample holders with a thin layer of carbon tape. Images were acquired using a Quanta 250 environmental scanning electron microscope (Thermo Scientific) in the high vacuum mode. The measurement of cell length was performed using ImageJ software (NIH).

### Auxin-induced growth

The shoot apexes and roots of 3-day-old *Arabidopsis* etiolated seedlings were removed using a razor blade. The hypocotyl segments were placed in the depletion medium (10 mM KCl, 1 mM MES, pH 6.0). Then the hypocotyl segments were transferred to new depletion medium with 1 μM IAA or 0.02% ethanol solvent control. Images of hypocotyl segments were acquired at the beginning of incubation and after 150 min of incubation (Fendrych *et al*., 2016).

### Fusicoccin-induced growth

The shoot apexes and roots of 3-day-old *Arabidopsis* etiolated seedlings were removed using a razor blade. The hypocotyl segments were placed in the depletion medium (10 mM KCl, 1 mM MES, pH 6.0). Then the hypocotyl segments were transferred to new depletion medium with 2 μM FC or 0.02% ethanol solvent control. Images of hypocotyl segments were acquired at the beginning of incubation and after 150 min of incubation.

### Creep Assay

The 3-day-old *Arabidopsis* etiolated seedlings were collected, stored at −80 °C, and thawed at room temperature prior to experiments. The seedlings were clamped under a constant force of 2.5 g. The clamped seedlings were incubated in 20 mM HEPES buffer (pH 6.8) for 20 min. Then the incubation buffer was replaced with 20 mM sodium acetate buffer (pH 4.5). The wall extension was recorded every 30 s for 75 min (Cosgrove, 1998; Durachko and Cosgrove, 2009).

For enzyme-induced creep, leaf petioles from the fifth to eighth leaves from 3-week-old plants were harvested, stored at −80 °C, and thawed prior to experiments. The petioles were abraded by rubbing the surface 10 times with carborundum slurry to facilitate buffer and protein penetration, and then washed with distilled water. The samples were pressed under a weight for 5 min to remove residual cell fluids and to flatten the wall sample, as described previously (Park and Cosgrove, 2012a, b). The pressed wall specimens were heat inactivated in boiling water for 12 s, and immediately cool down with cold running water. The wall specimens were clamped under a constant force of 8.2 g and incubated in 20 mM sodium acetate buffer (pH 4.5), which was replaced by the same buffer supplemented with 50 μg/mL GH5 family endoglucanase at the time of 20 min.

### Oryzalin Treatment

Wild type seeds were grown in the dark for 3 days on half-strength MS plate without sucrose and supplemented with 300 nM oryzalin. Oryzalin was dissolved in dimethyl sulfoxide (DMSO) to create stock solutions.

### Apoplastic pH Measurement

Seeds were grown on half-strength MS plates without sucrose for 4 days in the dark. A group of 30 seedlings were incubated in the depletion medium with 1 μM IAA or 0.02% ethanol solvent control. The pH of the incubation medium was measured using a pH meter before and after 150 min of incubation (Cleland, 1973).

The quantification of ratiometric pH sensor apo-pHusion in wild type and *csi1* was performed as described previously (Fendrych *et al*., 2016).

## Acknowledgement

We thank E. Wagner and X. Wang for their assistance with the creep assay. We thank D. Ehrhardt for providing YFP-CESA6 mCherry-TUA5 transgenic seeds. We thank A.T. Fuglsang for providing apo-pHusion transgenic seeds. This work was supported by the Center for LignoCellulose Structure and Formation, an Energy Frontier Research Center funded by the Department of Energy, Office of Science, Basic Energy Sciences under Award DESC0001090.

## Supplementary Figures

**Supplementary Fig. S1. A cartoon of the cross-section of an *Arabidopsis* hypocotyl.**

The outer epidermal cell wall is the periclinal wall that faces the external environment of the plant. The corresponding wall visualized by atomic force microscopy (AFM) and transmission electron microscopy (TEM) is shown. AFM examines the organization of the most recently deposited cellulose microfibrils in outer periclinal walls in *Arabidopsis* epidermal cells. TEM displays the structure of the entire outer epidermal wall. The white double-headed arrow on the AFM image indicates the long axis of the cell. The letter “c” on the TEM image indicates cytosol of the cell.

**Supplementary Fig. S2. Morphology of the *csi1-3* and *jia1-1* mutants.**

(A) Four-day-old dark-grown seedlings of wild type (Col-0), *csi1-3*, and *jia1-1* mutants (Scale bars, 5 mm).

(B) Ten-day-old dark-grown seedlings of wild type (Col-0), *csi1-3*, and *jia1-1* mutants (Scale bars, 5 mm).

(C) Measurement of cell length of epidermal cells in wild type (Col-0) etiolated seedlings grown for 2 days, 3 days, and 4 days.

(D) Measurement of cell length of epidermal cells in a file from the base to the top of 4-day-old, dark-grown hypocotyls of wild type (Col-0), *csi1-3*, and *jia1-1*.

**Supplementary Fig. S3. Auxin- and FC-induced hypocotyl elongation.**

(A) Dose-response graph of IAA on the hypocotyl length of wild type (Col-0). The lengths of the hypocotyls were measured at 0 min and 150 min of each treatment. Increase in length was calculated as [(length at 150 min – length at 0 min)/ length at 0 min] × 100%. Mock treatment was defined as using 0.02% ethanol (solvent) instead of IAA. ** *P* ≤ 0.01, *** *P* ≤ 0.001 (n = 6 for each treatment). Statistical analysis was performed by two-tailed Student’s *t*-test.

(B) Dose-response graph of FC on the hypocotyl length of wild type (Col-0). The lengths of the hypocotyls were measured at 0 min and 150 min of each treatment. Increase in length was calculated as [(length at 150 min – length at 0 min)/ length at 0 min] x 100%. Mock treatment was defined as using 0.02% ethanol (solvent) instead of FC. * *P* ≤ 0.05, ** *P* ≤ 0.01 (n = 6 for each treatment). Statistical analysis was performed by two-tailed Student’s *t*-test.

**Supplementary Fig. S4. Measurement of hypocotyl elongation rate.**

(A) Hypocotyl growth rate of wild-type (Col-0), *csi1-3*, and *jia1-1* mutants from day 2 to day 3. Seedlings were grown in the dark on vertical half-strength Murashige and Skoog (MS) plates without sucrose. The lengths of hypocotyls were measured at day 2 and day 3. Error bars represent SEM. *** *P* ≤ 0.001 (n =15 for each genotype). Statistical analysis was performed by two-tailed Student’s *t*-test.

(B) Auxin-induced hypocotyl elongation rate of wild-type (Col-0), *csi1-3*, and *jia1-1* mutants. The lengths of the hypocotyls were measured at 0 min and 150 min of 1 μM IAA treatment. Error bars represent SEM. *** *P* ≤ 0.001 (n = 12 for each genotype). Statistical analysis was performed by two-tailed Student’s *t*-test.

**Supplementary Fig. S5. CSC trajectories and cortical MTs are uncoupled in *csi1-3* mutant.**

(A-C) Two-channel confocal imaging of epidermal cells in 3-d-old etiolated seedlings from control and *csi1-3* mutant expressing YFP-CESA6 and mCherry-TUA5 in cell #17-21 (A), #9-13 (B), and #1-5 (C), respectively. Scale bars, 10 µm.

(D) Analysis of the average trajectory orientation of YFP-CESA6 and mCherry-TUA5 in control and *csi1-3* mutant in cell #17-21. Error bars are SEM. In control, the angles of YFP-CESA6 and mCherry-TUA5 trajectories are 17.87 ± 12.38 (mean ± SD) degrees and 19.67 ± 14.00 (mean ± SD) degrees, respectively. *P* > 0.05 (n = 6 for each group). In the *csi1-3* mutant, the angles of YFP-CESA6 and mCherry-TUA5 trajectories are 4.49 ± 4.00 (mean ± SD) degrees and 6.79 ± 2.49 (mean ± SD) degrees, respectively. *P* > 0.05 (n = 7 for each group).

(E) Analysis of the average trajectory orientation of YFP-CESA6 and mCherry-TUA5 in control and *csi1-3* mutant in cell #9-13. Error bars are SEM. In control, the angles of YFP-CESA6 and mCherry-TUA5 trajectories are 62.84 ± 17.27 (mean ± SD) degrees and 61.22 ± 13.35 (mean ± SD) degrees, respectively. *P* > 0.05 (n = 7 for each group). In the *csi1-3* mutant, the angles of YFP-CESA6 and mCherry-TUA5 trajectories are 11.07 ± 4.82 (mean ± SD) degrees and 57.79 ± 21.40 (mean ± SD) degrees, respectively. *P* ≤ 0.01 (n = 7 for each group).

(F) Analysis of the average trajectory orientation of YFP-CESA6 and mCherry-TUA5 in control and *csi1-3* mutant in cell #1-5. Error bars are SEM. In control, the angles of YFP-CESA6 and mCherry-TUA5 trajectories are 74.29 ± 8.13 (mean ± SD) degrees and 72.95 ± 8.91 (mean ± SD) degrees, respectively. *P* > 0.05 (n = 6 for each group). In the *csi1-3* mutant, the angles of YFP-CESA6 and mCherry-TUA5 trajectories are 21.82 ± 9.09 (mean ± SD) degrees and 77.69 ± 5.22 (mean ± SD) degrees, respectively. *P* ≤ 0.001 (n = 8 for each group).

## Notes

#### Summary of Updates

Supplemental files updated

