## Supplementary figures and images for "CELLULOSE SYNTHASE INTERACTIVE1- and Microtubule-Dependent Cell Wall Architecture Is Required for Acid Growth in *Arabidopsis Hypocotyls*"

### Figure S1

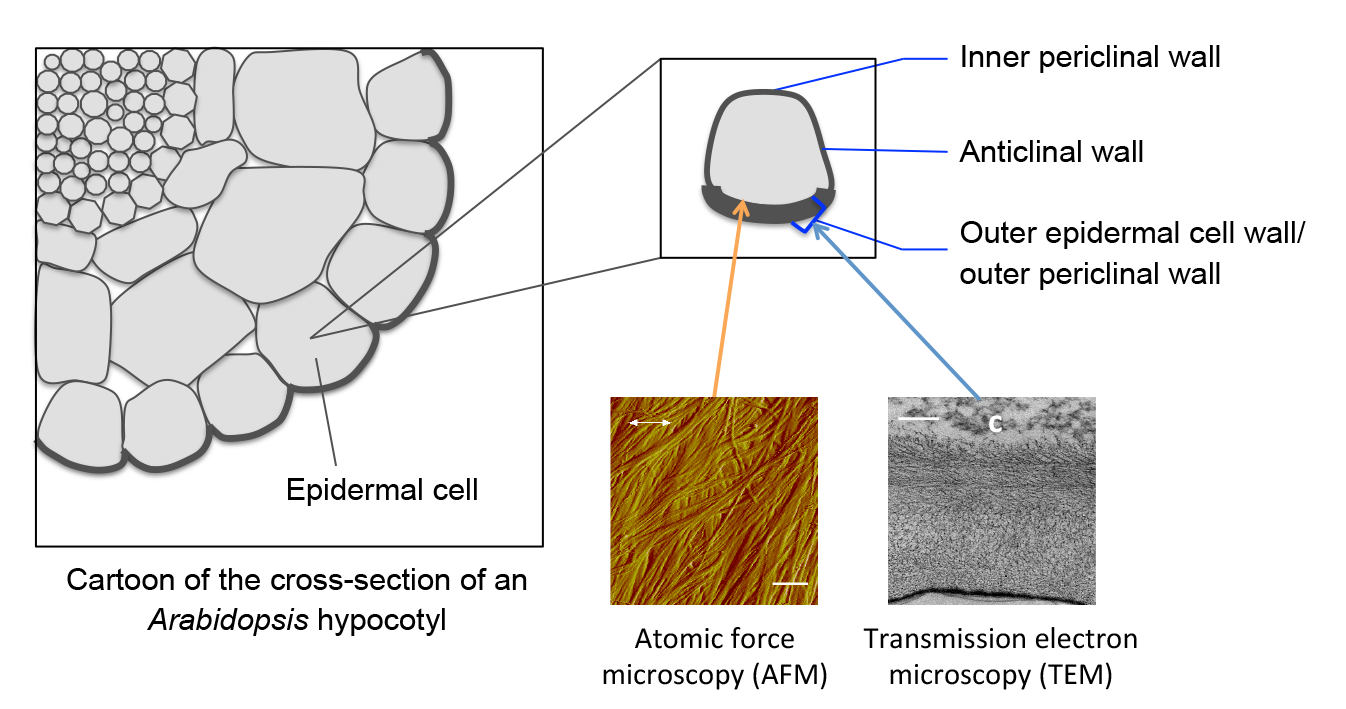

### Figure S2

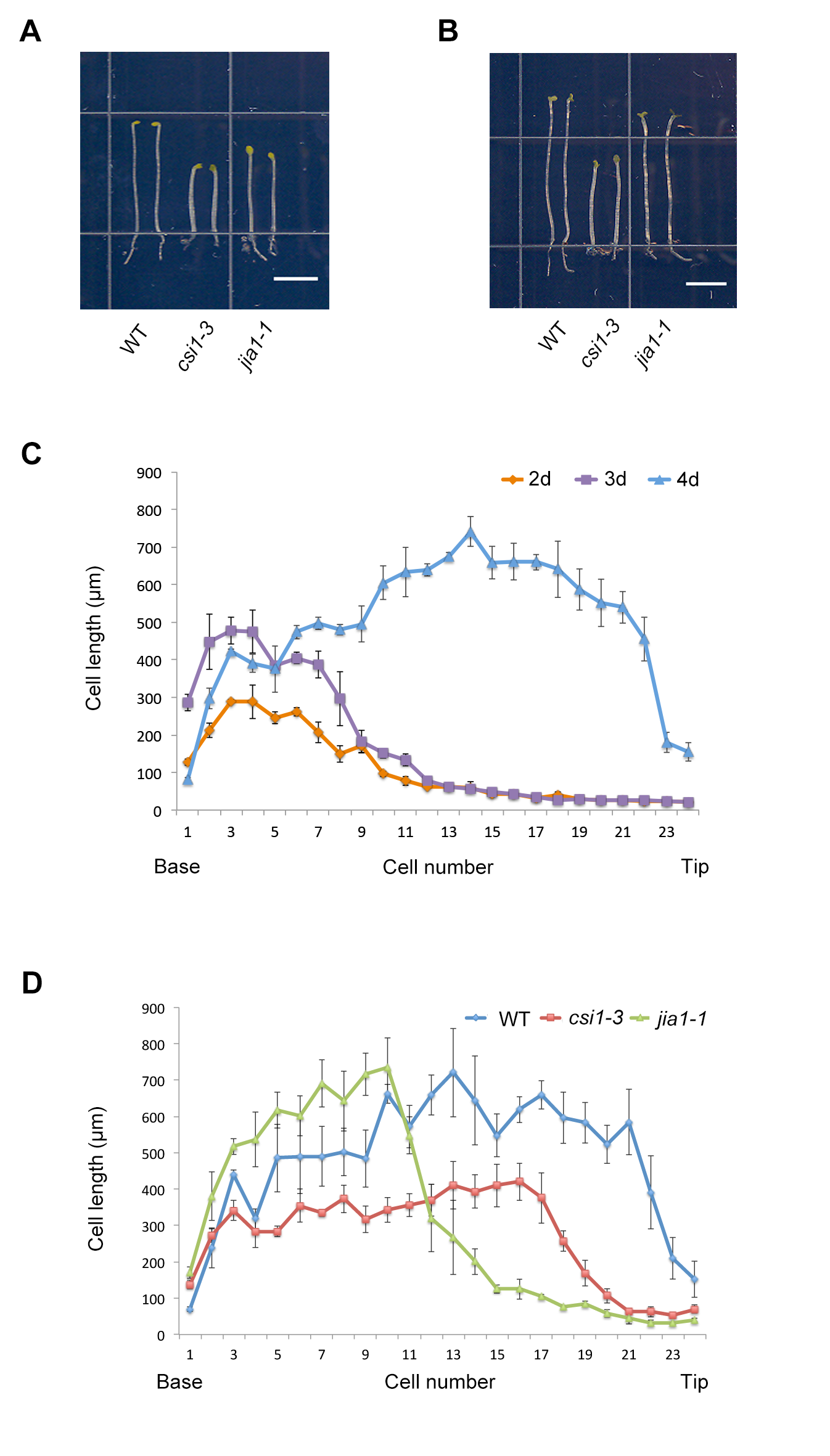

### Figure S3

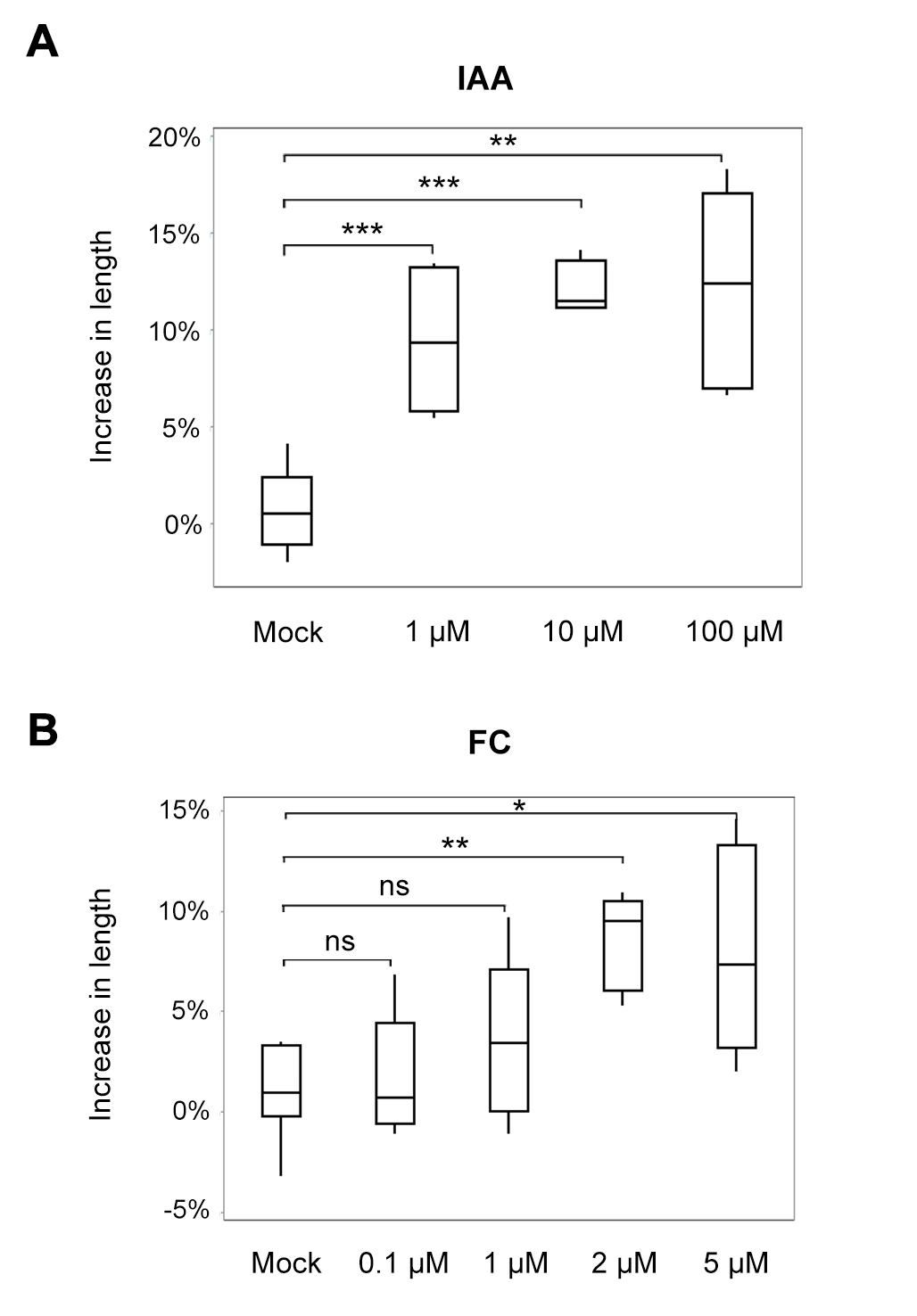

### Figure S4

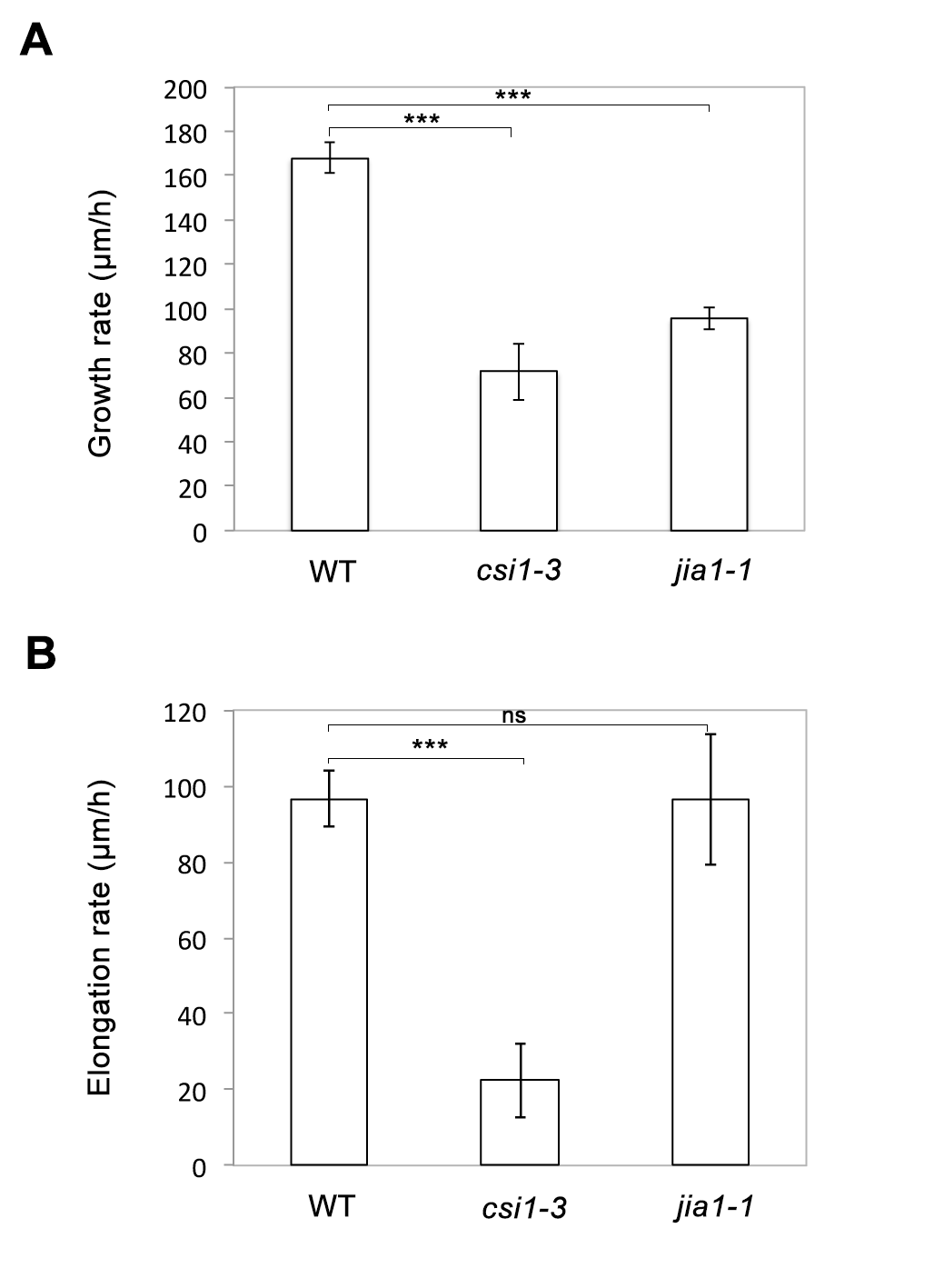

### Figure S5

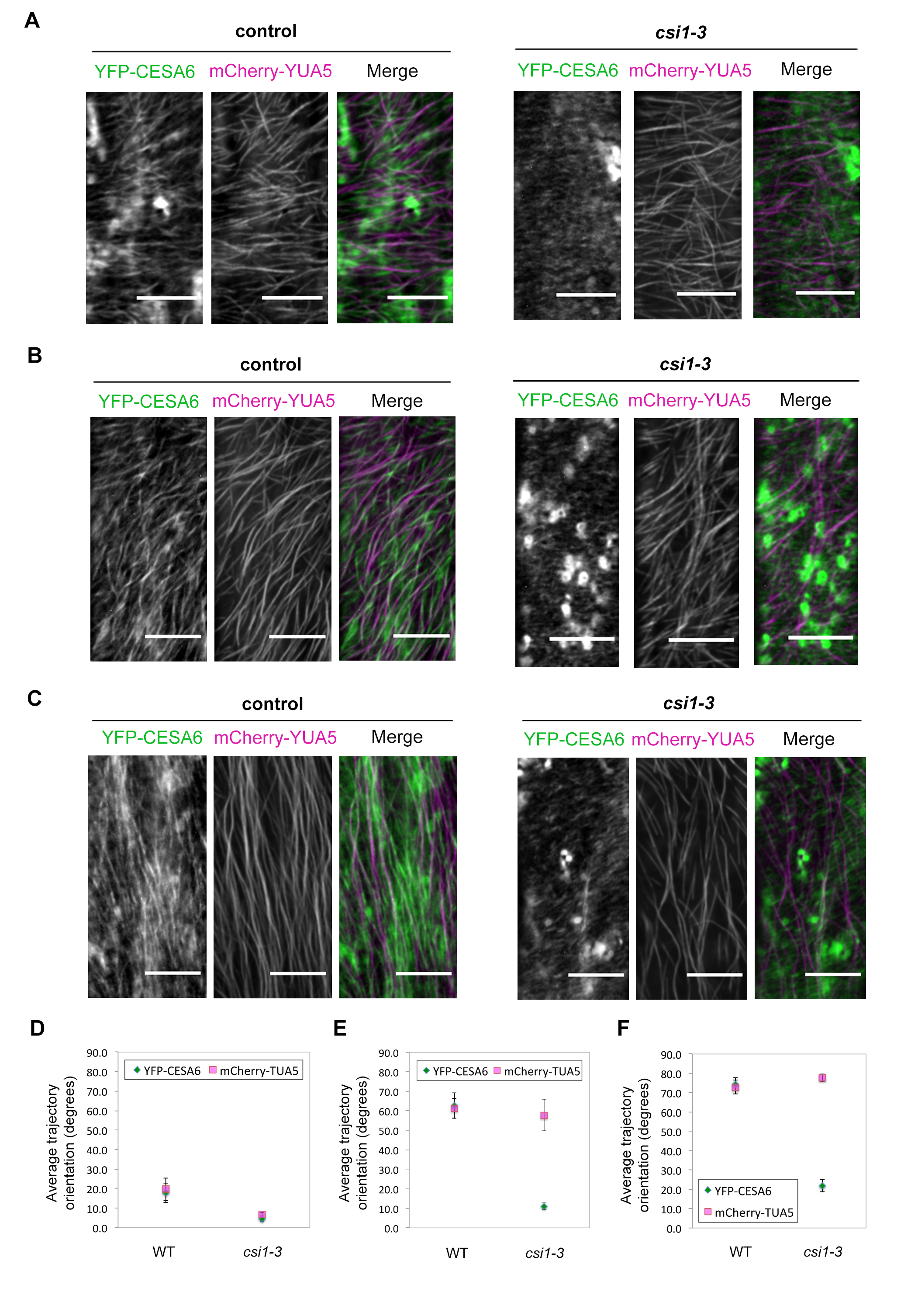
